# Three criteria of intrinsically theoretical categories in biological system and classification of some medical plants

**DOI:** 10.1101/814871

**Authors:** Huabin Zou

## Abstract

In classical biology, different taxonomic categories are all decided based on empirical rules established on learning knowledge. In taxonomy, different classification systems are of diversified rules. One biology is often in different categories, although the concepts related to taxonomic categories are the same to each other. Whether there exist some absolute standards to classify biology, and to give unalterable results. This is of greatly scientific meaning. Common and variation, that is heredity and variation are the most elemental information in biology, and exist at multiply biology material levels. In this paper a generalized biological heredity and variation information theory was proposed based on previous works. Three typical heredity and variation models were analyzed by using this theory. They are unique asymmetric variation model, symmetric two variation model and extreme radial variation model. In the maximum information states, two biological constants *P*_g1_ = 0.69 and *P*_g2_=0.61 and a boundary similarity function were obtained. These *P*_g_ and *P*_g_ function can be defined as the three theoretically taxonomic category criteria of biological system. For 29 samples belonging to four kind plants, their chemical fingerprint — infrared (IR) fingerprint spectra (FPS) were analyzed depending on the theoretical criteria. The correct classification ratio was 96.6%. The results showed these samples could be ideally classified. A suggestion was proposed that biology should be absolutely classified relying on the three intrinsic theoretical criteria.

## Introduction

Presently, taxonomy is a far from finished basically scientific research. All taxonomic categories are the classification grades in modern biology. These categories include Species, Genus (Genus), family (familia), Order (Ordo), Class (Classis), Phylum (Divisio), Kingdom (Regnum), which are all empirical rules determined relying on learning knowledge. In these rules, there is short of rigidly quantitative standards. Moreover, it lacks the support of mathematical principles. Currently, we can ask a question whether these categories really exist in biology, or whether there are some theoretical categories or grades in biology system. If some theoretical categories are deduced from some mathematical theories, whether they can correspond to taxonomic categories obtained by empirical knowledge, and whether these theoretical categories can be confirmed by experiments.

As we well known, biological common/heredity and variation are the most elemental information of biology, and exist at many material levels, including both biologically small molecules, macro-molecules, molecular structure, cell, organelle, organ, and individual, population, species and other higher levels, including Genus (Genus),family (familia),Order (Ordo), Class (Classis), Phylum (Divisio), Kingdom (Regnum). These can be named generally biological heredity and variation information. Whether a theory can be built up to describe generally biological heredity and variation information, to reveal some elemental laws and some particular laws, so as to classify biology accurately. This is a core theoretical problem in biological science. Research on common/heredity and variation information is the theoretical base of taxonomical science.

More than a century, the most important heredity and variation theory is population genetics [1-4] established grounded on Mondel laws and Hardy-Wenberger law at gene level in DNA sequence of the same biological species. In population genetics, the combination patterns or genotypes of allele and their distribution frequencies are investigated, and then their accompanying traits are researched. The effects of multiple genes on a biological characters are investigated too [5-9]. However, there is a short of study on heredity and variation laws at single base and base segments levels in DNA sequences so far.

Presently, statistical genetics and bioinformatics are the major theories to analyze heredity and variation at biological macro-molecule level besides population genetics. In bioinformatics, various variations are analyzed by sequence alignment, through comparing differences of structure units, such as single base in DNA, RNA sequences, various sequence segments in different genes, gene pools, and amino acid in protein sequences[8,10-15]. Grounded on above analysis, different kind phylogenetic trees of different biological samples, such as genes, proteins, organisms in different populations and species, can be achieved. Then taxonomy, classification, identification and cluster of biological individuals, genes, proteins are performed empirically[16-25]. In these methods, only the difference information are accepted to form empirical theories. While these methods are not able to reach a unchangeable results, which can outline the accurate laws, and do not vary with samples. Biological laws should be determined by common property of biology, so it is impossible to achieve the accurate law merely depending on variations.

For a long time, in biological science, there is no any accurate theory to deal with both common/heredity and variation simultaneously at any material level. Recently, it is necessary to built up the heredity and variation theory grounded on multiple material levels, such as molecule, cell, organ, individual and population, in order to describe unalterable heredity and variation laws.

How can we build up a theory to discover the intrinsic laws of common and variation. As well known, biological system is a physical chemistry system composed of thousands of substances. For a physical chemistry system, its physical chemistry action is in relatively steady states with certain action characteristics, which determine biological characteristics objectively. In modern biological science, biological chemistry reactions are investigated completely based on physical chemistry at molecular levels, and some simple movement rules are revealed at cell level[26-29].

On the other hand, as we well known, Shannon’s information theory suits to describe random system, but can not precisely represent a nonrandom system, such as biological system.

Theoretically speaking, a correctly general heredity and variation information theory should be grounded on physical chemistry action of heredity and variation substances, that is common heredity action and variation action. Author ZOU has deeply investigated this subject and proposed a theory, biological common heredity and variation information equation based on simple physical chemistry action model of heredity and variation substances [30]. Relying on the previous research fundamental [30,31,32,33,34,35], In this paper a generalized biological common heredity and variation information equation was further built up, theoretically which is qualified to describe heredity and variation at multiply material levels, such as various material structure units, biological molecules.

Based on the generalized biological common/heredity and variation information theory, three kind typical heredity and variation types were analyzed. Two biological constants and a boundary similarity function were obtained, which correspond to some biological categories. They can be defined as the theoretically taxonomic categories, and are tested by experiments in this research. The infrared fingerprint spectra, a kind of chemical fingerprint spectra of 29 samples of four kind medical plants were measured and analyzed. They could be divided into four classes ideally depending on these criteria, and their close relatives were correctly analyzed too.

The study showed these theoretical categories may correspond to some species, or to some genus, or other empirical categories. The boundary similarity function can be easily used as a rigid standard to determine which plants belong to close relative.

## 1 To establish generalized biological heredity and variation information theory

Heredity/common and variation wildly exist at many material levels, and result in generating and extincting of many biological systems. Because of the extreme complexity in biodiversity, which has a vast of variables, factors, the development in heredity and variation theory was limited greatly.

Whether there is a general theory to describe them? Author ZOU once investigated this subject based on physical chemistry principle systematically. Firstly, he independently defined and constructed the common peak (element, molecule) ratio *P*_g_, and variation peak (element, molecule) ratio *P*_vi_ scientifically in biological classification field [36,37,38,39]. Then dual index grade sequence individual pattern recognition method was proposed [40,41], based on the two index. These researches proved that the two index *P*_g_ and *P*_vi_ are able to represent biological properties well. Grounding on these investigations, a novel theory was proposed to deal with common heredity and variation systematically.

### 1.1 Generalized biological heredity and variation information theory

Depending on common peak (element, molecule) ratio *P*_g_, and variation peak (element, molecule) ratio *P*_vi_, author ZOU first proposed common heredity and variation information theory, that is dual index information theory, or heredity and variation information theory in 2009 [30]. The equation was as follows. When there are two kind variations *a, b*, corresponding to two samples in a biological system,

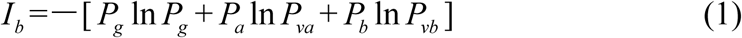

Where

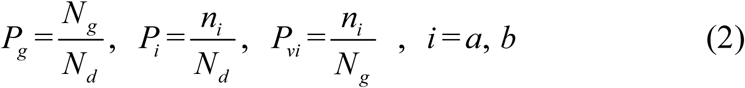

Generally, in the analysis of a sample set, one sample corresponds to one class of variation. If there are many kind variations in a biological system with a sample set, how to describe the action information. So a generalized equation need to be built up. Suggest there are *m* kind variations in a biological system, and each kind variation contain *n*_*i*_ elements or variables, then the generalized biological heredity and variation information equation was represented as follows.

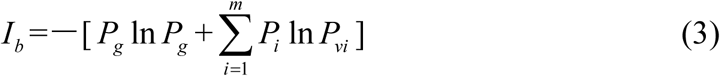

Where

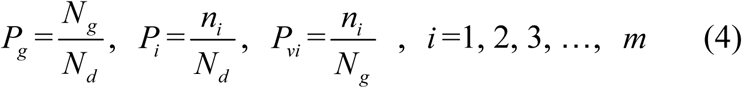

The meanings of variables in the equation (3), (4), as in [30,31,32,33,34,35] were listed bellow.

*P*_*g*_, common (heredity) peak(element, molecule)ratio, it can be briefly expressed as *P.*

This index is the same as the JaCcard and Sneath, Sokal coefficients.

*P*_*i*_, the ratio of *n*_*i*_ to *N*_*d*_. *P*_*vi*_ is the variation peak(element, molecule)ratio of *n*_*i*_ to *N*_*g*_.

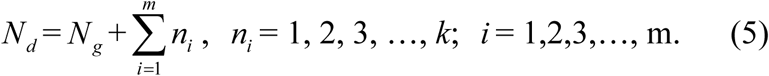

*N*_*d*_, the independent peaks (elements, molecules) existed in a system.

*N*_*g*_, common(heredity) peaks (elements, molecules) existed in a system.

*n*_*i*_, variation peaks (elements, molecules) belong to variation class *i*.

Relying on paper [30], — *P*_*g*_ ln *P*_*g*_ represents the similarity action of the system based on common elements, and 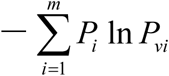represents the action between common and variation elements. According to these generalized biological heredity and variation information equation, a biological system is usually of three typical variation models. They are listed bellow.

Considering multiple types of variation elements, there are three typical variation models.

**Variation Model 1**, *n*_*i*_, *i* =1, *n*_1_>o, with only one variation class, the extreme asymmetric variation, or unique variation state in a system.

**Variation Model 2**, *n*_*i*_, *i* =1,2, and *n*_1_ = *n*_2_ >0, with two variation classes, or two symmetric variation states in a system.

**Variation Model 3**, *n*_*i*_ = 1, *i* = 1, 2, 3, …, *m*, with *m* classes of extreme radial variations. It means for each variation class, there is only one variation peak(element, molecule). This schematic was showed in figure 1.

**Fig. 1.**
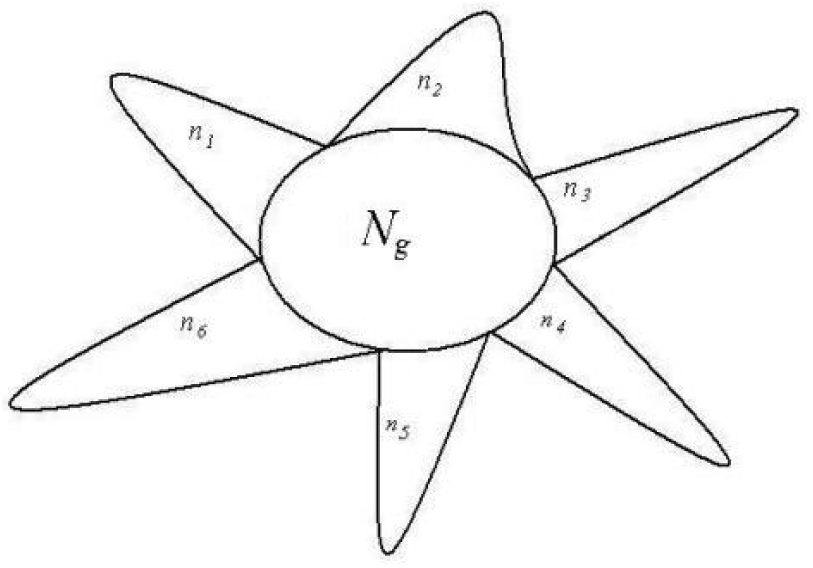
A system composed of *N*_g_ common heredity elements and *n*_*i*_ variation elements, *i* =1, 2, 3, …, *m*. For example, This system poses 6 classes of variation elements.

A lot of experiments have proved that for equation (1), it is able to accurately reveal some significant properties in complex biological systems. Two similarity constants *P*_g1_=0.69, *P*_g2_=0.61, can be obtained in the maximum information states. *P*_g1_=0.69 corresponds to the unique variation state, that is variation model 1. *P*_g2_=0.61 corresponds to symmetric variation state, that is variation model 2. These two similarity constants can be proved in **section1.2**. Based on the two similarity constants, very complex biological systems — the combination herbal medicines (traditional Chinese medicine, TCM), which consist of extracts of many medical plants, could be classified accurately relying on IR fingerprint spectra, which reflect structure unit information of biological small molecules [30,31,32,35]. Genus and Subgenus of pine could be discriminated precisely depending on information of molecular species in oleoresins [33], and four species of combination herbal medicines [33,35] were identified perfectly by means of the two constants. Three kinds of soybean proteomes were pattern recognized subtly by using the constant *P*_g2_=0.61, together with structure information of macro—molecules [34].

All these previous researches express that the primarily established heredity and variation information equation is qualified to uncover some heredity and variation laws of biology.

### 1.2 Similarity constants corresponding to Variation Model 1and Variation Model 2

#### Similarity constant *P*_g1_

When there is only single variation class, that is extreme asymmetric variation state, corresponding Variation Model 1. *n*_*i*_ >0, *i* = 1, *n*_1_ = *N*_*d*_ — *N*_*g*_, generally biological heredity and variation information equation can be expressed as follows.

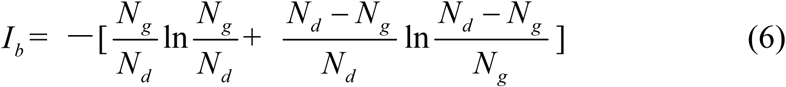

Let *N*_*d*_ = *N, N*_*g*_ = *x, I*_*b*_ = *y* (*x*), to substitute these into equation (6), get

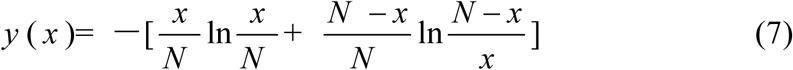

To take the derivative of equation (7), and let *I*_*b*_ to be in the maximum information state, then get

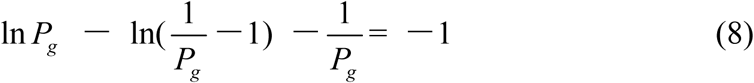

This equation has a solution *P*_g_=0.692, approximately is 0.69 = 69%. This theoretical standard has been successfully testified in researches [32,33,35]. *P*_g_=0.69 can be defined as similarity constant *P*_g1_.

According to article[30,31,32,33,34,35], for any two samples in a sample set, we can view them is a biological system. Theoretically, when their *P*_g_ ≧ 69%, they are in the asymmetric variation state, and they are of the identical properties. They belong to the same class. When their *P*_g_ < 69%, there are some distinct differences between them, they are not of identical property.

#### Similarity constant *P*_g2_

In a biological system, when in symmetric mutation state, corresponding to Variation Model 2,

*n*_1_ = *n*_2_ >0.

Let *n*_1_ = *n*_2_ = *n, n*_1_ + *n*_2_ = 2 *n* = *N*_*d*_ — *N*_*g*_, *n* > 0,

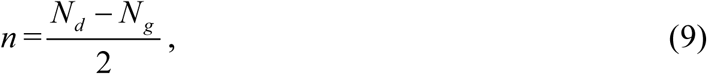

To substitute this formula into equation (3), then it becomes to be,

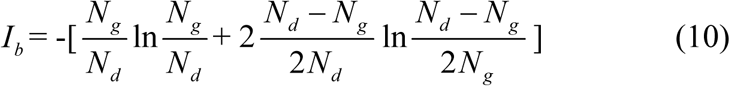

Let *N*_*d*_ = *N, N*_*g*_ = *x, I*_*b*_ = *y* (*x*) bring these to equation (10), then get

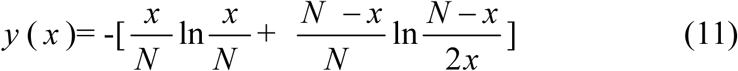

To take the derivative of equation (11), and order in the maximum information state, then get

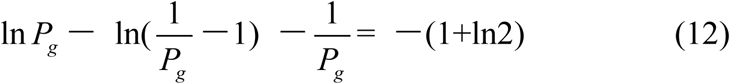

To solve this equation, one can obtain *P*_g_ = 0.6085, approximately is 0.61 =61%. *P*_g_ = 61% can be defined as similarity constant *P*_g2_.

According to article [30,31,32,33,34,35], for any two samples in a sample set, we can view them is a biological system. When their *P*_g_ ≧ 61%, they are of the identical properties, and they belong to the same class. when their *P*_g_ < 61%, there are some distinct differences between them, they are not of identical property.

In the same way, one can get,

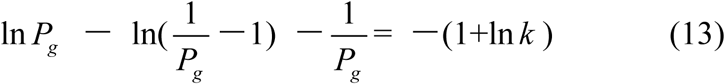

For *k* classes of variations.

### 1.3 Boundary Similarity function corresponding to Variation Model 3

For extreme radial variation model, *n*_*i*_ = 1, *i* = 1,2,3,…, *m*. that is all variation elements are different from one another. Each variation element belongs to its own class. In this situation, generally biological heredity and variation information equation can be represented as follows.

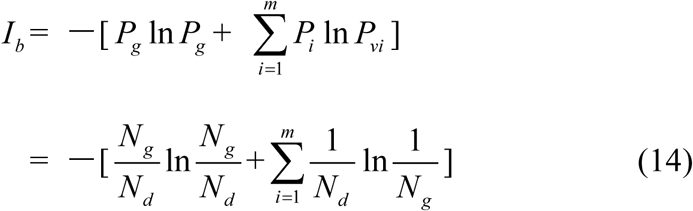

As before, let *m* = *N*_*d*_ — *N*_*g*_ = *N* — *N*_*g*_, and order *I*_*b*_ = y, *N*_*g*_ = *x, N*_*d*_ = *N*, to substitute them into equation(14).
then get

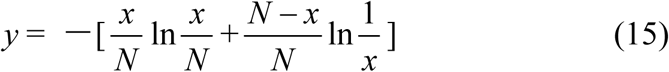

To take the derivative of equation (15), and let it to be in the maximum information state, then obtain,

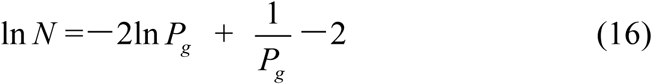

As *N*→ ∞, *P*_g_ <<1, then 1 *P*_*g*_ >> 2ln *P*_*g*_ equation (16) can be simplified to be

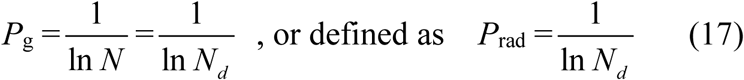

Considered some degree of randomness of heredity and variation in biology, we can deduce a formula for accurately and briefly calculating the common peak (molecules, or any elements) ratio *P*_*g*_ in biology,

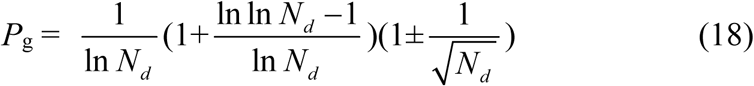

Since the theoretical criteria are similarity *P*_g_, then these *P*_g_ show the ratios of common materials to total materials. The common characteristics reflect the identical properties, which are the ancestor characters of these samples in a biological system. So the value of *P*_g_ reflects high or low of ancestor traits between these samples. The higher the *P*_g_, the closer the relative relationship of them. So to determine relative relationship far and near between samples relying on the high or low of *P*_g_ is scientifically reasonable.

Equation (17), (18) represent the relationship between *P*_g_ and *N*. That is there is no constant *P*_g_ in Variation Model 3. They are also the boundary condition or maximum variation range for close relatives, which originate from the same close ancestor. Since there is no limitation in the number and kinds of samples in a biological system, this theory is suitable for any open biological system.

The similarity function (17), (18) can be defined as the discrimination function of close relative biology. This function is the critical point for extreme variation of a biological system. For any two samples in a sample set, we can view them is a biological system. When their *P*_g_ ≧ *P*_rad_, they are of the certain identical properties, and they belong to close relative. When their *P*_g_ < *P*_rad_, there are extremely distinct differences between them, they do not belong to close relative. *P*_rad_ reflects the theoretical boundary of the close relative relationship.

Theoretically, these intrinsic criteria of *P*_g_ should be related to some biological ranks, that is theoretically taxonomic categories, such as species, genus, family and so on. Then these intrinsically characteristic *P*_g_ can be defined as differently theoretical criteria of some intrinsically taxonomic categories.

As we known, there are many biological levels or ranks, including species, genus, family as well as others in relative systems. Whether these ranks correspond to these *P*_g_ criteria for intrinsically taxonomic categories, or they can be theoretically discriminated by these criteria, previously classified by empirical knowledge. All these need to be verified by experiments.

Certainly, for the two constants, they may correspond to two intrinsically taxonomic categories, which should correspond to any two different biological ranks, such as species and subspecies, genus, subgenus, family and so on. These have been proved successfully by some researches [33,34]. These researches indicated the theoretically taxonomic categories, existed intrinsically in the equation, do not one— to — one correspond to empirical ranks of biology, obtained by means of empirical classification. From the view of scientific point, people should classify biology into some absolute ranks based on these theoretical criteria, and should offer strictly theoretical boundaries of close relatives.

Presently, The major methods in biological classification, cluster and pattern recognition are classical taxonomy[42-46], molecular taxonomy [16-25], Chemical taxonomy [42-44]. The major study manner is to discover new rules by means of experiments or depending on empirical knowledge, but not by some mathematical principles. On the other hand, many similarity and difference index were applied for the recognition, classification, cluster and evolution researches of traditional medical plants and combination herbal medicines, soybeans [30-32,34-41,47-52], depending on IR fingerprint spectra, one of the chemical fingerprint spectra, information of biological molecular structure units. This may be a good approach for building up new mathematical principles existed in biological ranks.

## 2 Classification of 29 medical plant samples based on three theoretical criteria

### 2.1 Plant samples

In this article IR fingerprint spectra of 29 plant samples were measured and analyzed, the experiments, seen **Methods** in section 4. these samples belong to four kind medical plants: Baishao, Chishao, Huangqi, and Gancao. In these samples, Baishao, Chishao belong to *Paeonia* L.(genus), Huangqi belongs to *Astragalus* Linn.(genus), Gancao belongs to *Glycyrrhiza* Linn.(genus). The sources of these 29 samples were listed in table 1.

**Table 1.** The sources of 24 herbal samples ^a^.

| Sample | Chinese name | Latin name | Originated district in China | Collecting time |
| --- | --- | --- | --- | --- |
| S1 | **Baishao <sup>b</sup>** | *Paeonia Lactiflora* Pall <sup>d</sup> | Bozhou of Anhui province | 2002.9 |
| S2 |  | *Paeonia Lactiflora* Pall | Bozhou of Anhui province | 2002.9 |
| S3 |  | *Paeonia Lactiflora* Pall | Xinji in Heze city of Shandong province | 2003.10 |
| S4 |  | *Paeonia Lactiflora* Pall | Huangti in Heze city of Shandong province | 2003.10 |
| S5 |  | *Paeonia Lactiflora* Pall | Houji in Heze city of Shandong province | 2003.10 |
| S6 |  | *Paeonia Lactiflora* Pall | Bozhou of Anhui province | 2003.11 |
| S7 |  | *Paeonia Lactiflora* Pall | Hangzhou city of Zhejiang province | 2003.10 |
| S8 | **Chishao <sup>c</sup>** | *Paeonia Lactiflora* Pall | Jilin province | 2007.5. |
| S9 |  | *Paeonia Lactiflora* Pall | Inner mongolia | 2006. |
| S10 |  | *Paeonia Lactiflora* Pall | Henan province | 2006. |
| S11 |  | *Paeonia Lactiflora* Pall | Liaoning province | 2006. |
| S12 |  | *Paeonia Lactiflora* Pall | Shandong province | 2006. |
| S13 |  | *Paeonia Lactiflora* Pall | Jinan city of Shandong province | 2007. |
| S14 |  | *Paeonia Lactiflora* Pall | Jianlian drug store of Jinan | 2004.2 |
| S15 | **Huangqi** | Astragalus membranaceus (Fisch.)Bge. | Inner mongolia | 2007.2 |
| S16 |  | Astragalus membranaceus (Fisch.)Bge. | Inner mongolia | 2006 |
| S17 |  | Astragalus membranaceus (Fisch.)Bge. | Inner mongolia | 2006 |
| S18 |  | Astragalus membranaceus (Fisch.)Bge. | Gansu province | 2006 |
| S19 |  | Astragalus membranaceus (Fisch.)Bge. | Longxi of Gansu province | 2006 |
| S20 |  | Astragalus membranaceus (Fisch.) | Jilin province | 2007.10 |
| S21 | **Gancao** | Glycyrrhiza uralensis fisch | Hangjinhou qi of Bayannaoer meng in inner Mongolia | 2001.10 |
| S22 |  | Glycyrrhiza uralensis fisch | Yikezhao meng of inner Mongolia | 2001.10 |
| S23 |  | Glycyrrhiza uralensis fisch | Inner mongolia | 2002.3 |
| S24 |  | Glycyrrhiza uralensis fisch | Inner mongolia | 2002.9 |
| S25 |  | Glycyrrhiza uralensis fisch | Left qi of Alashan meng of inner Mongolia | 2002.10 |
| S26 |  | Glycyrrhiza uralensis fisch | Right qi of Alashan meng of inner Mongolia | 2002.10 |
| S27 |  | Glycyrrhiza uralensis fisch | Ejina qi of Alashan meng of inner Mongolia | 2002.10 |
| S28 |  | Glycyrrhiza inflata Batal | Shihezi city of Xinjiang | 2002.12 |
| S29 |  | Glycyrrhiza uralensis fisch | Yongdeng of Gansu province | 2002.10 |

*a.* All samples were the dried roots of these medical plants. *b.* Baishao is the cultured *Paeonia Lactiflora* Pall in traditional Chinese medicine. *c.* Chishao is the wild *Paeonia Lactiflora* Pall. *d.* All samples were identified by the chief pharmacist Kang Shuang-long, Yao Ting-zhi and by Professor Zhou Feng-qin.

All samples were kept at bellow —18°C, after collected and dried.

### 2.2 Analysis on IR fingerprint spectra data of 29 plant samples

#### 2.2.1 Determination and represent of common and variation peaks in IR fingerprint spectra

According to the methods described in section 4, to extract major compositions of samples with absolute ethanol, and to measure their IR fingerprint spectra. Then to determine the common and variation peaks of the these IR FPS by means of Shapiro-Wilk test [53]. The common and variation peaks were listed in table 2.

**Table 2.** Codes of absorption peaks in IR fingerprint spectra of 29 samples.

| Sample | Codes of absorption peaks |
| --- | --- |
| S1 | 1, 4, 8<sup>a</sup>, 13, 17, 21, 26, 29, 34, 37, 43, 52, 56 |
| S2 | 3, 8, 13, 17, 21, 26, 29, 34, 37, 43, 52, 56 |
| S3 | 3, 8, 13, 14, 16, 21, 26, 29, 34, 37, 39, 52, 54, 56, 59 |
| S4 | 3, 8, 13, 16, 21, 26, 29, 34, 37, 43, 52, 57, 59 |
| S5 | 3, 8, 13, 16, 21, 26, 29, 33, 39, 43, 52, 57 |
| S6 | 3, 8, 13, 16, 21, 26, 29, 34, 37, 39, 43, 47, 52, 57, 59 |
| S7 | 4, 8, 13, 16, 21, 26, 29, 34, 39, 44, 47, 52, 57 |
| S8 | 5, 7, 8, 13, 17, 21, 26, 29, 40, 52, 60 |
| S9 | 5, 7, 8, 13, 17, 21, 26, 29, 39, 46, 50, 52, 55, 57 |
| S10 | 5, 7, 8, 13, 15, 17, 21, 26, 29, 39, 46, 50, 52, 55, 57, 59 |
| S11 | 5, 7, 8, 13, 17, 21, 26, 29, 39, 46, 50, 52, 55 |
| S12 | 5, 8, 13, 17, 21, 26, 29, 38, 39, 50, 52, 55, 58 |
| S13 | 5, 8, 13, 17, 21, 26, 29, 38, 39, 50, 52, 55, 57 |
| S14 | 1, 2, 8, 13, 16, 21, 26, 29, 39, 50, 52, 54, 57, 59 |
| S15 | 6, 8, 12, 16, 18, 21, 23, 38, 41, 48, 53, 60 |
| S16 | 6, 8, 12, 16, 18, 23, 38, 41, 48, 53, 60 |
| S17 | 6, 8, 12, 16, 23, 24, 28, 41, 53, 60 |
| S18 | 6, 8, 12, 16, 23, 31, 39, 41, 60 |
| S19 | 6, 8, 12, 16, 23, 24, 41, 60 |
| S20 | 6, 8, 12, 16, 23, 24, 31, 38, 41, 53, 60 |
| S21 | 1, 8, 9, 17, 19, 20, 23, 27, 29, 32, 35, 39, 42, 44, 49, 59 |
| S22 | 1, 8, 15, 19, 20, 23, 27, 32, 35, 39, 42, 44, 49, 58 |
| S23 | 1, 8, 10, 17, 19, 20, 23, 27, 32, 35, 39, 42, 44, 49, 59 |
| S24 | 4, 8, 9, 17, 19, 20, 22, 25, 27, 29, 32, 36, 37, 42, 44, 49, 58 |
| S25 | 4, 8, 9, 17, 19, 20, 23, 27, 32, 36, 37, 42, 44, 49, 58 |
| S26 | 4, 8, 9, 17, 19, 20, 23, 24, 27, 29, 32, 37, 44, 45, 49, 51, 58 |
| S27 | 1, 8, 11, 17, 19, 20, 23, 27, 32, 37, 44, 49, 58 |
| S28 | 2, 8, 9, 17, 19, 20, 23, 24, 32, 36, 37, 42, 44, 49, 59 |
| S29 | 2, 8, 15, 19, 23, 27, 30, 36, 39, 42, 44, 49, 57 |

$\alpha$ , 1,4,8 means the first, fourth, eighth peak in IR fingerprint spectra of 29 plants. Other codes are of the same meaning.

The averaged wavenumbers (cm^-1)^ of all peaks are listed bellow.

1,3417±6; 2,3406±1; 3,3396±2; 4,3385±2; 5,3375±3; 6,3363±2; 7,2973±1; 8,2928±2; 9,2113±2; 10, 2102±1; 11,2090±1; 12,1727±3; 13,1714±3; 14,1642± 1; 15,1635±1; 16,1624±2; 17,1615±2; 18,1570±1; 19,1512±1; 20,1458±1; 21,1450±2; 22,1419±1; 23,1408±4; 24,1385±4; 25,1372±1; 26,1346±3; 27,1333±1; 28,1300±1; 29,1278±3; 30,1259±1; 31,1252±1; 32,1234±1; 33,1227±1; 34,1204±1; 35,1131±3; 36,1103±2; 37,1073±2; 38,1062±1; 39,1051±2; 40,1037±1; 41,1029±1; 42,997±1; 43,938±3; 44,926±2; 45,892±1; 46,880±1; 47,871±1; 48,856±1; 49,835±1; 50,821±1; 51,777±1; 52,714±1; 53,667±3; 54,632±2; 55,623±3; 56,613±1; 57,596±5; 58,582±2; 59,553±4; 60,526±1. Peaks with the same code in different samples belong to a common peak group. There are 60 common peak groups or independent peaks in the 29 plant samples.

#### 2.2.2 Characteristic sequences and classification of 29 plant samples

According to the two similarity constants *P*_g1_ = 69%, *P*_g2_ = 61%, obtained from generally biological heredity and variation information equation (3), and the analysis in [30,31,32,33,34,35], for any two samples in a biological system, when the *P*_g_ scale is *P*_g_ ≧(61±3) %, these two samples are similar to each other greatly and of the same inner quality. The similarities among these 29 samples were showed **in table 3.1, 3.2.** In table 3.1 and 3.2, the same kind/class plants were showed in the same color area. From these tables, the similarities among the same kind plant samples were significantly higher than that among different kind plant samples. Common peak ratios among different kind plant samples are much less than *P*_g2_ constant 61%.

**Table 3.1.** Common peak ratios among the 29 samples and their cluster results

| Sample | S1 | S2 | S3 | S4 | S5 | S6 | S7 | S8 | S9 | S10 | S11 | S12 | S13 | S14 | S15 | S16 | S17 | S18 | S19 | S20 | S21 | S22 | S23 | S24 | S25 | S26 | S27 | S28 | S29 |
| --- | --- | --- | --- | --- | --- | --- | --- | --- | --- | --- | --- | --- | --- | --- | --- | --- | --- | --- | --- | --- | --- | --- | --- | --- | --- | --- | --- | --- | --- |
| S1 | **100<sup>a</sup>** | **79<sup>b</sup>** | 47 | 50 | 39 | 56 | 44 | 41 | 35 | 32 | 37 | 37 | 37 | 35 | 9 | 4 | 5 | 5 | 5 | 4 | 16 | 8 | 8 | 20 | 17 | 20 | 18 | 12 | 4 |
| S2 | **79** | **100** | **59<sup>c</sup>** | **63** | 50 | 50 | 39 | 44 | 37 | 33 | 39 | 39 | 39 | 30 | 9 | 5 | 5 | 5 | 5 | 5 | 12 | 4 | 8 | 16 | 5 | 16 | 14 | 13 | 4 |
| S3 | 47 | **59** | **100** | **61** | 50 | 58 | 47 | 30 | 32 | 35 | 33 | 33 | 33 | 53 | 13 | 8 | 9 | 14 | 10 | 8 | 15 | 7 | 11 | 10 | 7 | 10 | 8 | 11 | 8 |
| S4 | 50 | **63** | **61** | **100** | **63** | **71** | 50 | 32 | 33 | 36 | 29 | 29 | 35 | 47 | 13 | 9 | 9 | 10 | 10 | 9 | 11 | 4 | 7 | 11 | 7 | 11 | 8 | 12 | 8 |
| S5 | 39 | 50 | 50 | **63** | **100** | **59** | 56 | 35 | 44 | 40 | 39 | 39 | 47 | 53 | 14 | 10 | 10 | 17 | 11 | 10 | 12 | 8 | 8 | 7 | 4 | 7 | 4 | 4 | 14 |
| S6 | 56 | 50 | 58 | **71** | **59** | **100** | **65** | 30 | 38 | 41 | 33 | 33 | 40 | **61** | 13 | 8 | 9 | 14 | 10 | 8 | 19 | 12 | 11 | 10 | 7 | 10 | 12 | 11 | 12 |
| S7 | 44 | 39 | 47 | 50 | 56 | **65** | **100** | 33 | 42 | 38 | 37 | 37 | 44 | 50 | 14 | 9 | 10 | 16 | 11 | 9 | 16 | 13 | 12 | 15 | 12 | 15 | 8 | 8 | 18 |
| S8 | 41 | 44 | 30 | 32 | 35 | 30 | 33 | **100** | 56 | 50 | **60** | 50 | 50 | 32 | 15 | 10 | 11 | 11 | 12 | 10 | 13 | 4 | 8 | 12 | 8 | 12 | 9 | 8 | 4 |
| S9 | 35 | 37 | 32 | 33 | 44 | 38 | 42 | 56 | **100** | **88** | **93** | **69** | **80** | 47 | 8 | 4 | 4 | 10 | 5 | 4 | 15 | 8 | 12 | 11 | 7 | 11 | 8 | 7 | 13 |
| S10 | 32 | 33 | 35 | 36 | 40 | 41 | 38 | 50 | **88** | **100** | **81** | **61** | **71** | 50 | 8 | 4 | 4 | 9 | 4 | 4 | 19 | 11 | 15 | 10 | 7 | 10 | 7 | 11 | 16 |
| S11 | 37 | 39 | 33 | 29 | 39 | 33 | 37 | **60** | **93** | **81** | **100** | **73** | **73** | 42 | 9 | 4 | 5 | 10 | 5 | 4 | 16 | 8 | 12 | 11 | 8 | 11 | 8 | 8 | 8 |
| S12 | 37 | 39 | 33 | 29 | 39 | 33 | 37 | 50 | **69** | **61** | **73** | **100** | **86** | 42 | 14 | 9 | 5 | 10 | 5 | 9 | 16 | 13 | 12 | 15 | 12 | 15 | 13 | 8 | 8 |
| S13 | 37 | 39 | 33 | 35 | 47 | 40 | 44 | 50 | **80** | **71** | **73** | **86** | **100** | 50 | 14 | 9 | 5 | 10 | 5 | 9 | 16 | 8 | 12 | 11 | 8 | 11 | 8 | 8 | 13 |
| S14 | 35 | 30 | 53 | 47 | 53 | **61** | 50 | 32 | 47 | 50 | 42 | 42 | 50 | **100** | 13 | 9 | 9 | 15 | 10 | 9 | 20 | 12 | 16 | 7 | 4 | 7 | 8 | 12 | 17 |

*a.* the common peak ratio of sample its own is 100% . *b.* common peak ratio $P_g$ of S1 and S2 is 79%. *c.* common peak ratio $P_g \geq 59\%$ , expressed by boldface in different color areas.

**Table 3.2.** Common peak ratios among the 29 samples and their cluster results

| Sample | S1 | S2 | S3 | S4 | S5 | S6 | S7 | S8 | S9 | S10 | S11 | S12 | S13 | S14 | S15 | S16 | S17 | S18 | S19 | S20 | S21 | S22 | S23 | S24 | S25 | S26 | S27 | S28 | S29 |
| --- | --- | --- | --- | --- | --- | --- | --- | --- | --- | --- | --- | --- | --- | --- | --- | --- | --- | --- | --- | --- | --- | --- | --- | --- | --- | --- | --- | --- | --- |
| S15 | 9 | 9 | 13 | 13 | 14 | 13 | 14 | 15 | 8 | 8 | 9 | 14 | 14 | 13 | 100 | 92 | 57 | 50 | 54 | 64 | 8 | 8 | 8 | 4 | 8 | 7 | 9 | 8 | 9 |
| S16 | 4 | 5 | 8 | 9 | 10 | 8 | 9 | 10 | 4 | 4 | 4 | 9 | 9 | 9 | 92 | 100 | 62 | 54 | 59 | 69 | 8 | 9 | 8 | 4 | 8 | 8 | 9 | 8 | 9 |
| S17 | 5 | 5 | 9 | 9 | 10 | 9 | 10 | 11 | 4 | 4 | 5 | 5 | 5 | 9 | 57 | 62 | 100 | 58 | 80 | 75 | 8 | 9 | 9 | 4 | 9 | 13 | 10 | 14 | 10 |
| S18 | 5 | 5 | 14 | 10 | 17 | 14 | 16 | 11 | 10 | 9 | 10 | 10 | 10 | 15 | 50 | 54 | 58 | 100 | 74 | 67 | 14 | 15 | 14 | 4 | 9 | 8 | 10 | 9 | 16 |
| S19 | 5 | 5 | 10 | 10 | 11 | 10 | 11 | 12 | 5 | 4 | 5 | 5 | 5 | 10 | 54 | 59 | 80 | 74 | 100 | 73 | 9 | 10 | 10 | 4 | 10 | 14 | 11 | 15 | 11 |
| S20 | 4 | 5 | 8 | 9 | 10 | 8 | 9 | 10 | 4 | 4 | 4 | 9 | 9 | 9 | 64 | 69 | 75 | 67 | 73 | 100 | 8 | 9 | 8 | 4 | 8 | 12 | 9 | 13 | 9 |
| S21 | 16 | 12 | 15 | 11 | 12 | 19 | 16 | 13 | 15 | 19 | 16 | 16 | 16 | 20 | 8 | 8 | 8 | 14 | 9 | 8 | 100 | 67 | 72 | 50 | 55 | 50 | 53 | 55 | 45 |
| S22 | 8 | 4 | 7 | 4 | 8 | 12 | 13 | 4 | 8 | 11 | 8 | 13 | 8 | 12 | 8 | 9 | 9 | 15 | 10 | 9 | 67 | 100 | 61 | 41 | 53 | 41 | 59 | 38 | 59 |
| S23 | 8 | 8 | 11 | 7 | 8 | 11 | 12 | 8 | 12 | 15 | 12 | 12 | 12 | 16 | 8 | 8 | 9 | 14 | 10 | 8 | 72 | 61 | 100 | 39 | 50 | 39 | 47 | 58 | 56 |
| S24 | 20 | 16 | 10 | 11 | 7 | 10 | 15 | 12 | 11 | 10 | 11 | 15 | 11 | 7 | 4 | 4 | 4 | 4 | 4 | 4 | 50 | 41 | 39 | 100 | 78 | 62 | 50 | 52 | 25 |
| S25 | 17 | 5 | 7 | 7 | 4 | 7 | 12 | 8 | 7 | 7 | 8 | 12 | 8 | 4 | 8 | 8 | 9 | 9 | 10 | 8 | 55 | 53 | 50 | 78 | 100 | 68 | 65 | 67 | 33 |
| S26 | 20 | 16 | 10 | 11 | 7 | 10 | 15 | 12 | 11 | 10 | 11 | 15 | 11 | 7 | 7 | 8 | 13 | 8 | 14 | 12 | 50 | 41 | 39 | 62 | 68 | 100 | 58 | 52 | 25 |
| S27 | 18 | 14 | 8 | 8 | 4 | 12 | 8 | 9 | 8 | 7 | 8 | 13 | 8 | 8 | 9 | 9 | 10 | 10 | 11 | 9 | 53 | 59 | 47 | 50 | 65 | 58 | 100 | 47 | 30 |
| S28 | 12 | 13 | 11 | 12 | 4 | 11 | 8 | 8 | 7 | 11 | 8 | 8 | 8 | 12 | 8 | 8 | 14 | 9 | 15 | 13 | 55 | 38 | 58 | 52 | 67 | 52 | 47 | 100 | 33 |
| S29 | 4 | 4 | 8 | 8 | 14 | 12 | 18 | 4 | 13 | 16 | 8 | 8 | 13 | 17 | 9 | 9 | 10 | 16 | 11 | 9 | 45 | 59 | 56 | 25 | 33 | 25 | 30 | 33 | 100 |

The definition of characteristic sequence:

For sample S*i*, its characteristic sequence is composed of these samples, whose common peak ratios related to S*i* fit to *P*_g_ scale *P*_g_ ≧(61±3) %. According to table 3.1, and 3.2, the characteristic sequences of 29 samples were obtained and displayed in table 4. Their classification were performed based on their characteristic sequences.

**Table 4.** Characteristic sequences of 29 plant samples.

| Class | Sample | Characteristic sequence | Class | sample | Characteristic sequence |
| --- | --- | --- | --- | --- | --- |
| **Baishao<sup>a</sup>** | **S1** | S1 S2 | **Huangqi<sup>c</sup>** | **S15** | S15 S16 S20 |
|  | **S2** | S1 S2 S3 S4 |  | **S16** | S15 S16 S17 S19 S20 |
|  | **S3** | S2 S3 S4 |  | **S17** | S16 S17 S19 S20 |
|  | **S4** | S2 S3 S4 S5 S6 |  | **S18** | S18 S19 S20 |
|  | **S5** | S4 S5 S6 |  | **S19** | S16 S17 S18 S19 S20 |
|  | **S6** | S4 S5 S6 S7 |  | **S20** | S15 S16 S17 S18 S19 S20 |
|  | **S7** | S6 S7 | **Gancao<sup>d</sup>** | **S21** | S21 S22 S23 |
| **Chishao<sup>b</sup>** | **S8** | S8 S11 |  | **S22** | S21 S22 S23 S27 |
|  | **S9** | S9 S10 S11 S12 S13 |  | **S23** | S21 S22 S23 |
|  | **S10** | S9 S10 S11 S12 S13 |  | **S24** | S24 S25 S26 |
|  | **S11** | S8 S9 S10 S11 S12 S13 |  | **S25** | S24 S25 S26 S27 S28 |
|  | **S12** | S9 S10 S11 S12 S13 |  | **S26** | S24 S25 S26 |
|  | **S13** | S9 S10 S11 S12 S13 |  | **S27** | S22 S25 S27 |
|  | **S14** | S6 S14 |  | **S28** | S25 S28 |
|  |  |  |  | **S29** | S22 S29 |

*a.* Baishao is *Paeonia Lactiflora* Pall (cultured), *b.* Chishao is *Paeonia Lactiflora* Pall (wild), *c.* Huangqi belongs to *Astragalus* Linn., *d.* Gancao belongs to *Glycyrrhiza* Linn.

In accordance with the characteristic sequences of these 29 samples, the characteristic sequences in one class of samples were made up of themselves. Samples in one kind samples’ characteristic sequences form an independent sample set. There is no overlapped sample among different sample sets, expect for S14, whose characteristic sequence contains S6. In the 29 samples, 28 samples were classified correctly by means of their characteristic sequences. The correct ratio is 28/29 = 96.6%. Relying on the theoretical standard *P*_g_ scale *P*_g_ ≧(61±3) %, of the intrinsically taxonomic category, the four kind plants Baishao S1-S7, Chishao S8-S13, Huangqi S15-S20, Gancao S21-S29, were accurately classified, or pattern recognized.

In particular, the significantly similar samples themselves in characteristic sequences are the best markers of their classes. This property is very important for classification, identification and pattern recognition of samples.

Summery, the similarities among Baishao and Chishao samples were much higher than that among any other two classes, such as Huangqi and Gancao samples, Baishao and Huangqi samples. It indicated that Baishao and Chishao samples should be close relatives.

These above results showed that similarity constant *P*_g2_ = 61% can be used as the quantitative standard to discriminate which plants are of the same efficiency, or used as the standard to determine the identical herbal medicines [30,31,32,35].

On the other hand, suggesting *m* = 4, that is there are four classes of plants, the theoretical standard *P*_g_ scale is *P*_g_ ≧53.2%, relying on the equation (13). To analyze the data in table 3.1 and 3.2, and get characteristic sequences of these samples. The results were very similar to that obtained by means of the standard *P*_g_ scale *P*_g_ ≧(61±3) %. There were trivial differences in their characteristic sequences. This also proved that generally biological heredity and variation information equation can represent intrinsic properties of biology exactly.

#### 2.2.3 Analysis on close relatives among the four class plants

If these four kind plants are in the maximum variation state, that is at extreme radial variation state, accurate *P*_g_ scale *P*_g_≧24.4% (0.269=26.9%, according to equation (18)) could be obtained, when *N* = 60, by equation (17). *P*_g_ scale *P*_g_≧24.4% is the theoretical boundary of close relatives of these 29 plant samples. Relying on this standard to determine characteristic sequences of these 29 samples, they could be classified into three classes perfectly, showed as follows.

**Characteristic sequences of these 29 samples A class: Baishao and Chishao (***Paeonia Lactiflora* Pall )

S1:S1 S2 S3 S4 S5 S6 S7 S8 S9 S10 S11 S12 S13 S14^a^

S2:S1 S2 S3 S4 S5 S6 S7 S8 S9 S10 S11 S12 S13 S14

S3:S1 S2 S3 S4 S5 S6 S7 S8 S9 S10 S11 S12 S13 S14

S4:S1 S2 S3 S4 S5 S6 S7 S8 S9 S10 S11 S12 S13 S14

S5:S1 S2 S3 S4 S5 S6 S7 S8 S9 S10 S11 S12 S13 S14

S6:S1 S2 S3 S4 S5 S6 S7 S8 S9 S10 S11 S12 S13 S14

S7:S1 S2 S3 S4 S5 S6 S7 S8 S9 S10 S11 S12 S13 S14

S8:S1 S2 S3 S4 S5 S6 S7 S8 S9 S10 S11 S12 S13 S14

S9:S1 S2 S3 S4 S5 S6 S7 S8 S9 S10 S11 S12 S13 S14

S10:S1 S2 S3 S4 S5 S6 S7 S8 S9 S10 S11 S12 S13 S14

S11:S1 S2 S3 S4 S5 S6 S7 S8 S9 S10 S11 S12 S13 S14

S12:S1 S2 S3 S4 S5 S6 S7 S8 S9 S10 S11 S12 S13 S14

S13:S1 S2 S3 S4 S5 S6 S7 S8 S9 S10 S11 S12 S13 S14

S14:S1 S2 S3 S4 S5 S6 S7 S8 S9 S10 S11 S12 S13 S14

**B class: Huangqi (***Astragalus* Linn.)

S15:S15 S16 S17 S18 S19 S20

S16:S15 S16 S17 S18 S19 S20

S17:S15 S16 S17 S18 S19 S20

S18:S15 S16 S17 S18 S19 S20

S19:S15 S16 S17 S18 S19 S20

S20:S15 S16 S17 S18 S19 S20

**C class: Gancao (***Glycyrrhiza* Linn.)

S21: S21 S22 S23 S24 S25 S26 S27 S28 S29

S22: S21 S22 S23 S24 S25 S26 S27 S28 S29

S23: S21 S22 S23 S24 S25 S26 S27 S28 S29

S24: S21 S22 S23 S24 S25 S26 S27 S28

S25: S21 S22 S23 S24 S25 S26 S27 S28 S29

S26: S21 S22 S23 S24 S25 S26 S27 S28

S27: S21 S22 S23 S24 S25 S26 S27 S28 S29

S28: S21 S22 S23 S24 S25 S26 S27 S28 S29

S29: S21 S22 S23 S25 S27 S28 S29

*a*. S1:S1 S2 S3 S4 S5 S6 S7 S8 S9 S10 S11 S12 S13 S14, represents the characteristic sequence of sample S1. Other sequences are of the same meaning.

From their characteristic sequences showed in the above classification results, these four kind plants could be divided into three close relatives. A class: Baishao +Chishao (*Paeonia Lactiflora* Pall). B class: Huangqi (*Astragalus* Linn.), C class: Gancao (*Glycyrrhiza* Linn.). This conclusion comply with the empirical classification. While for the classification based on medical efficiency and the kind of herbal medicines, theoretical standard *P*_g_ scale *P*_g_≧(61±3) % is more reasonable, see table 4.

These results also indicated that for species, genus, they could be also clustered excellently by means of the theoretical boundary function(17),(18). This theoretical boundary can tolerate larger randomness of similarity in the same species, genus and so on. This theoretical boundary also suits to express elemental characteristics of biological evolution. Especially, the results of theoretical classification and close relative analysis both represented that common peak ratio *P*_g2_ = 61%, indeed reflect the intrinsic characteristics of biological systems. Based on classical taxonomy, Baishao and Chishao both belong to *Paeonia* L.(genus), the same species, *Paeonia Lactiflora* Pall. However, there are some differences between them. Baishao is the plant *Paeonia Lactiflora* Pall having been cultured by human for thousand years. But Chishao is wild *Paeonia Lactiflora* Pall. Moreover, there exist great differences in their medical efficiency between Baishao and Chishao. This indicates their chemical compositions vary obviously. According to pharmacopoeia of P.R.China[54], two kinds of Huangqi *Astragalus membranaceus* (Fisch.)Bge. and *Astragalus membranaceus* (Fisch.), all belong to *Astragalus* Linn.(genus), their medical efficiency are identical. Two kind gancao: *Glycyrrhiza uralensis* fisch and *Glycyrrhiza inflata* Batal, they all belong to genus *Glycyrrhiza* Linn.(genus). Their medical efficiency are very similar to each other. The classification results relying on the theoretical standard *P*_g2_ scale *P*_g_ ≧(61±3) % proved these two kind Gancao were in the same class, so were the two kind Huangqi medicine plants.

Baishao and Chishao are viewed as the same species, but their chemical compositions and medical efficiency are all different from each other distinctly. Theoretically, they should belong to different species, depending on researches in this article.

Huangqi and Gancao belong to the same family Leguminosae, but different genus. Two kind Huangqi samples belong to two species, and the same genus. In terms of the theoretical standard *P*_g_ scale *P*_g_ ≧(61±3) %, Huangqi samples are in the same class. So are the Gancao samples. These indicated for *Glycyrrhiza* Linn. and *Astragalus* Linn., *P*_g_ scale *P*_g_ ≧(61±3) % may be their theoretical standard of genus as well as that of herbal medicines, which are of the identical efficiency.

Depending on the above analysis, we know plant evolution is of some universal laws. Two similarity constants *P*_g1_, *P*_g2_, some times are suitable to be the theoretical standards of biological species, some time they are suitable for the theoretical standard of genus. Thus an hypothesis that biological system can be classified into some intrinsically taxonomic categories based on the three intrinsically theoretical criteria, can be proposed. Contrast to traditional classification, or taxonomy, only relying on empirical knowledge, these approaches can not reach the accurate and unchangeable results.

Nowadays, an reasonable classification scheme should be that all biology are firstly classified into the three intrinsic grade patterns, then divided these samples into sub-patterns in terms of the classical taxonomy methods.

## 3 Conclusion on intrinsic taxonomic categories and their theoretical criteria

In plant systematics, there is no quantitative classification criteria for species, genus, and family, and other biological ranks [42,43,45,46]. The results of this article showed there are three theoretical criteria corresponding to three intrinsically taxonomic categories, which correspond to three typically biological Variation Models. For *P*_g_ scales, *P*_g_ ≧(61±3) %, *P*_g_ ≧(69±3) %, they can be accepted as the theoretical standards for some species, some genus, or some families. For *P*_g_ ≧ 1/ ln *N*_*d*_, it may be used as the theoretical standard to discriminate close relatives, such as Genus, Family (familia), Order (Ordo), Class (Classis), Phylumo (Divisi), Kingdom (Regnum) for some plants. This indirectly indicated that for different biological systems, their evolution speed vary significantly.

Most interestingly, these criteria may be a strong tool to investigate some laws in biologically macro molecules, such as DNA, RNA and protein sequences.

The research results also showed generally biological heredity and variation information theory, which was constructed based on the physical chemistry action model of heredity and variation materials, was able to accurately describe some heredity and variation laws in biological system, and suitable for many material levels. On the other hand, how to accurately determine the close relative among different biological systems is a very fundamental science subject. A theoretical boundary function 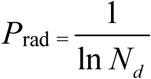, or is precisely equal to 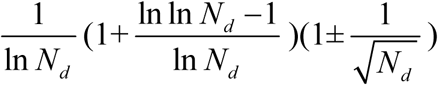 was derived from generally biological heredity and variation information equation. It can be used as a strong tool for determining close relatives.

The laws or rules of material actions are usually more universal and more rigorous than that obtained by statistics. Through actions of substances, which produce differently biological phenomena, the strictly, scientifically and universally theoretical standards for classification should be established. This progress may promote biological science to some extent in the future, and open the door towards quantitative researches on intrinsically scientific principles of biology.

## 4 Methods

### 4.1 Instrument

Vector 22 FT-IR (Bruker scientific technology Co., Ltd., Germany), spectra range:4 000 cm^-1^ — 400 cm^-1^, Resolution 4 cm^-1^. High speed mill, Analytical balance (METTLE TOLEDO), sensitivity 0.1 mg. Soxhlet extractor. Hot water bath.

### 4.2 Reagent

Chloroform (AR), ethanol (AR)(Kemiou chemical regent limited company, Tianjin, China).

### 4.3 Preparation of samples and measurement condition

To dry the root samples of 29 plants at 60°C for 2 hours, then to grind these dried roots into powders and sieve these powders with 60 mesh. 2.00 g powders (3 parallels for each sample)of one sample were taken and wrapped with filter paper, which was put into soxhlet extractor. Then 50 ml of chloroform was put into the extractor. To extract sample for 2 hours at boiling point of chloroform, and pour out the extracted solution, in order to extract little of liposoluble substances.

The residual chloroform solved in powders was evaporated clearly by using hair dryer with hot air for 5 minutes. Then to put 60 ml of absolute ethanol into Soxhlet extractor, and extract sample for 1.5 hours. The extraction was poured out into an evaporator on hot water bath at 80 °C, till the solvent was evaporated clearly.

After this above step, the dried extract was resolved by using dehydrated ethanol, treated with molecular sieve, to be a saturated solution. It was put into sample tube and kept at below 0 °C.

The IR FPS of samples were measured with liquid film method based on KBr crystal flat. Each extract was measured three times, and each sample with three parallel extracts, then 9 IR FPS were yield. To take the mean wavenumbers of each group of common peaks in these 9 IR FPS as a combination peak, all combination peaks of a sample form its combination IR FPS. Then all combination IR FPS of these 29 samples were compared to form multiple groups of common peaks, which were analyzed by means of the approach showed in this paper. The sensitivity of vector 22 FT-IR was 2. For extracts kept at 0∼ 4 °C, they were of good repeatability.

### 4.4 Software

All data were analyzed by means of the software written by my students.

## Acknowledgement

I sincerely thank academician WANG yongyan of Chinese Engineering Academy for his moral encouragement and support.

